# A trans-acting enhancer lncRNA modulates androgen-dependent gene expression via sequence-specific interaction with the Androgen Receptor

**DOI:** 10.1101/2025.05.09.652846

**Authors:** R. Sultanov, A. Suzdalenko, P. Shnaider, O. Zubkova, G. Arapidi

**Affiliations:** Lopukhin Federal Research and Clinical Center of Physical-Chemical Medicine of Federal Medical Biological Agency, Moscow, Russia; Shemyakin-Ovchinnikov Institute of Bioorganic Chemistry of the Russian Academy of Sciences, Moscow, Russia

## Abstract

**Background:** Enhancer RNAs (eRNAs) have been shown to modulate the transcriptional landscape of prostate cancer (PCa). The Androgen Receptor (AR), a well-known modulator of eRNA expression, contains an RNA-binding region that interacts with RNA molecules in a sequence-specific manner. However, there is currently no evidence that AR forms complexes with eRNAs to regulate gene expression.

**Methods:** To explore the eRNA-AR interactome, we reanalyzed publicly available RNA-seq, ChIP-seq, and AR-RIP-seq data from prostate cancer cell lines to identify a single long non-coding eRNA that interacts with AR (ARA-elncRNA1). Using linear regression, we identified genomic regions where AR occupancy is associated with the expression level of ARA-elncRNA1. We further demonstrated that this eRNA recruits AR to YY1-mediated enhancer-promoter loops, stabilizing these interactions. A series of experiments, including RIP-qPCR, ChIP-qPCR, and EMSA, were conducted to validate the proposed model. Finally, we investigated the role of ARA-elncRNA1 in prostate cancer cells survival.

**Results:** We confirmed the sequence-specific interaction between AR and ARA-elncRNA1. This eRNA not only regulates AR occupancy at the promoters of several AR-dependent genes but also protects AR from proteasomal degradation. The AR:ARA-elncRNA1 complex interacts with YY1 and influences enhancer-promoter looping. Additionally, ARA-elncRNA1 was found to inhibit the proliferation of prostate cancer (PCa) cells *in vitro* and expression level of lncRNA was associated with a lower Gleason score.

**Conclusions:** Our findings revealed the existence of an eRNA that directly binds to AR and regulates, *in trans*, the expression of several AR-dependent genes. We demonstrated that eRNAs can not only interact with AR but also modulate chromatin structure. These insights shed new light on the functional roles of eRNAs and their contribution to cancer development.

## Introduction

Long non-coding RNAs (lncRNAs) have long outgrown the initial contemptuous attitude toward them. We now understand their significant role in various cellular processes, ranging from transcription regulation to intercellular communication [1, 2]. Among the diverse types of lncRNAs, one of the most enigmatic is enhancer RNAs (eRNAs)—transcripts expressed from enhancer regions [3–6]. Most eRNAs are non-polyadenylated, unspliced, and considered transcriptional noise with no known functions [7]. However, some models propose that eRNAs facilitate enhancer-promoter contacts through mechanisms such as liquid-liquid phase separation [8].

A small fraction of eRNAs are stable, polyadenylated (~10%), and even spliced (~5%), resembling typical long non-coding RNAs [7]. For example, a 2-kb eRNA specific to muscle tissue contains an intron and plays a significant role in the expression of Myogenin by recruiting cohesin [9]. Another example is heart-specific enhancer lncRNAs, whose expression is linked to pathophysiological models of heart disease *in vivo* [10]. Despite these specific cases, the general functional role of most enhancer lncRNAs (elncRNAs) remains poorly understood.

The androgen receptor (AR) is a well-studied hormone receptor that plays a crucial role in prostate development and disease [11]. While the ligand-and DNA-binding properties of AR have been extensively characterized, its RNA-binding activity was only recently discovered [12–14]. Karyn Schmidt and colleagues identified an RNA-binding motif in the N-terminal domain of AR, specifically recognizing a 9-nucleotide sequence (CYUYUCCWS, where Y = pyrimidine [C or U], W = weak [A or U], and S = strong [C or G]) [13]. This discovery unveiled a new layer of transcriptional regulation mediated by AR. Subsequent studies have demonstrated interactions between AR and various lncRNAs, including *SRA1*, *HOTAIR*, *PCGEM1*, and the eRNA of the *KLK3* gene [15]. With this RNA motif identified, it is now possible to systematically investigate the spectrum of AR-binding RNAs and their potential roles in prostate development and disease.

Here, we report the identification of a novel member of the elncRNA family—AR-binding long non-coding enhancer RNA 1 (ARA-elncRNA1)—which trans-regulates the expression of several AR-dependent genes. We demonstrate that this eRNA influences AR recruitment to gene promoters lacking canonical androgen response elements (AREs) by forming a complex with the transcription factor YY1. Additionally, ARA-elncRNA1 protects AR from proteasomal degradation. Functional studies reveal that this eRNA plays a role in prostate cancer cell lines proliferation and colony formation. Notably, its expression negatively correlates with prostate cancer grade.

Our findings suggest that sequence-specific binding of AR to eRNAs represents a novel regulatory mechanism for controlling AR-dependent gene expression in a hormone-independent manner.

## Results

### Investigation of AR binding eRNAs

To conduct a comprehensive catalog of enhancer lncRNAs in LNCaP cells, we combined PolyA+ RNA-seq datasets generated by different research groups (Table S1) and used the Scallop2 [16] tool for *de novo* assembly of 31009 genes (File S1). We filtered out all annotated genes (GENCODE v37), genes with a high Ribo-seq signal in LNCaP cells (Table S1), genes shorter than 200 bp, and genes with an expression level below 1 TPM (Figure 1A). This procedure resulted in 1840 novel lncRNA genes (Table S2). To annotate transcripts as AR-binding, we utilized AR RIP-seq (RNA immunoprecipitation following sequencing) data from LNCaP cells [14] (Table S1, S2) and selected only transcripts containing the AR-binding RNA motif and enriched in RNA fraction precipitated with AR specific antibodies (logFC>0.5 and FDR< 0.05). Out of the 1840 transcripts, only 13 exhibited AR-binding properties (Figure 1A,B, Table S2).

**Figure 1.**
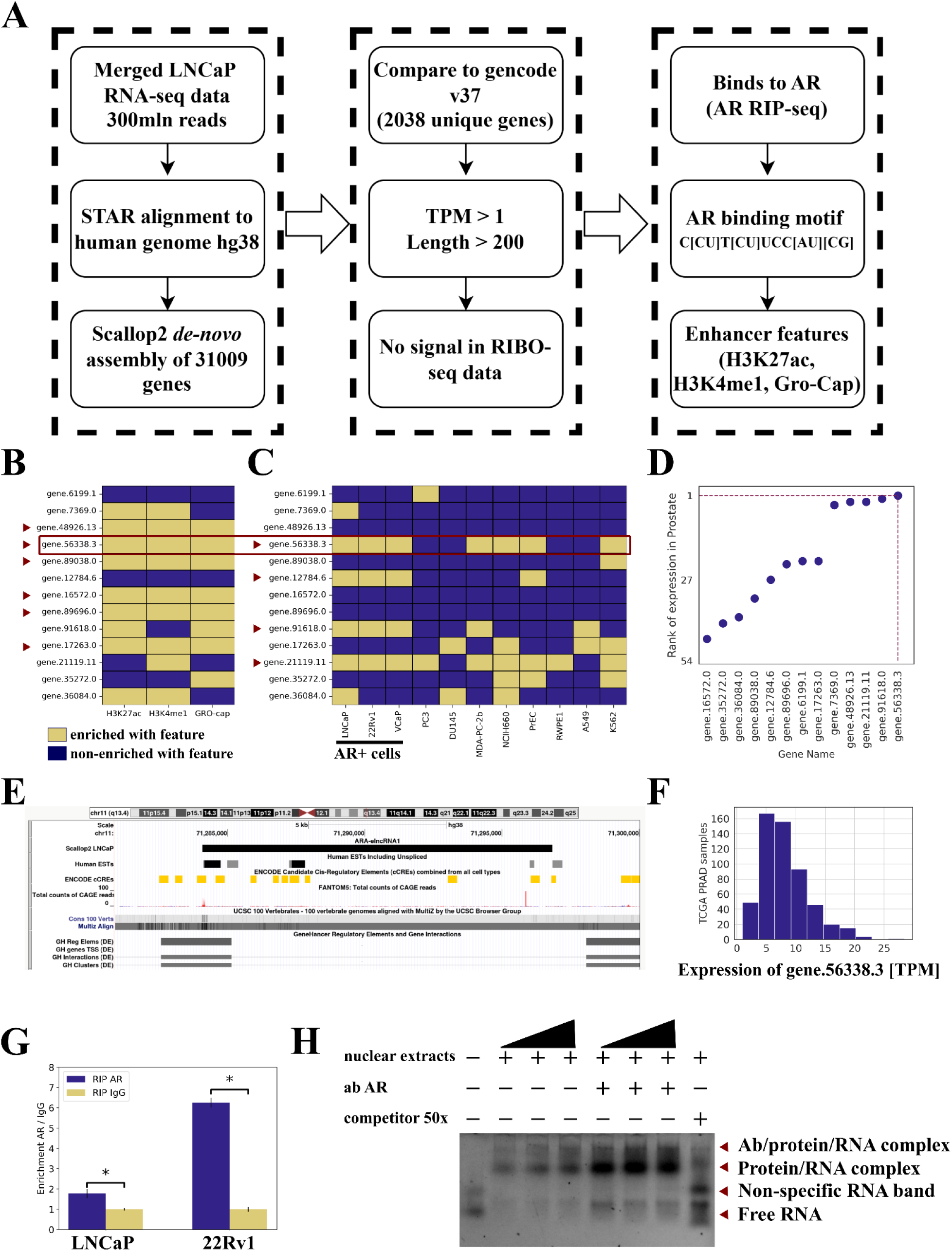
Identification of AR-binding eRNAs A. Schematic illustration of the computational pipeline used to establish a comprehensive catalog of eRNAs in the LNCaP cell line. B. Heatmap showing the enrichment of eRNA transcription start sites (TSS) by enhancer features. Red triangles indicate eRNAs enriched by all examined features. C. Heatmap of eRNA expression in selected CCLE cell lines. Red triangles indicate transcripts expressed in all three AR-positive cell lines (AR+ cells) with an expression level higher than 0.5 TPM. The red rectangle highlights the transcript that meets all filtration criteria. D. Dot plot displaying the rank of expression for each eRNA in prostate GTEx samples. The lowest rank corresponds to the highest expression level. E. Snapshot of the UCSC Genome Browser showing RefSeq gene annotation, EST data, ENCODE CRE track, 100-vertebrate genome conservation track, FANTOM5 CAGE-seq data track, and GeneHancer database track. F. Histogram showing the distribution of ARA-elncRNA1 expression levels in TCGA PRAD samples. G. RIP results confirming the interaction between ARA-elncRNA1 and AR in the nuclear fraction of LNCaP and 22Rv1 cells. RT-qPCR was used to examine enrichment of ARA-elncRNA1 in the anti-AR antibody fraction of RIP compared to IgG fraction. “*” - p-value<0.05 (two-tailed unpaired Student’s t-test). Results are presented as mean ± SEM calculated from three replicates. H. RNA electrophoretic mobility shift assay (EMSA). A Cy3-labeled RNA probe containing the AR-binding motif was incubated with increasing amounts of 22Rv1 nuclear extracts and resolved on a 1% TBE agarose gel. Specificity of AR binding was tested by adding anti-AR antibodies or an unlabeled competitor probe.

Next, we applied an “enhanceability” filter, selecting only transcripts with high levels of H3K27ac, H3K4me1, and PRO-Cap signals in LNCaP cells (Table S1, S2, Figure 1B, red triangles). We also examined the expression of these 13 transcripts in prostate cell lines from the CCLE (Cancer Cell Line Encyclopedia) [17] panel and retained those expressed in AR-positive cells (Figure 1C). Finally, we ranked all transcripts based on their expression in normal prostate tissue from the GTEx database [18] and filtered out those with low ranks corresponding to the highest expression level (Figure 1D, Figure S1A). Only one genomic region met all filtration criteria: gene.56338.3.

Using a UCSC genome browser [19], we visually confirmed that expression of the identified enhancer region is supported by CAGE-seq data from the FANTOM5 database (Figure 1E) [20]. The existence of this transcript was further validated by known expression sequence tags (EST database), and its transcription start site (TSS) region was found to be highly conserved among primates (Figure 1E). Additionally, the TSS region of gene.56338.3 contains a cis-regulatory elements (CREs) from the ENCODE database [21] and is identified as an enhancer for the *NUMA1* and *SHANK2* protein-coding genes in the GeneHancer database [22] (Figure 1E). Also this transcript is highly expressed in TCGA PRAD (The Cancer Genome Atlas, prostate adenocarcinoma) samples (Figure 1D). Collectively, these findings confirm that the transcript from the enhancer gene.56338.3 is an enhancer RNA (eRNA) that may bind to AR.

Based on Scallop2 *de novo* assembly, gene.56338.3 produces 10 polyexonic and 2 monoexonic transcripts. By inspecting RNA-seq datasets from LNCaP, 22Rv1, and VCaP cell lines (Table S1) in the IGV browser [23], we confirmed that the major form of these transcripts is the monoexonic transcript gene.56338.3.4, which is 4468 nt in length (Figure S1A, File S2). The other transcripts were expressed at negligible levels in the studied cell lines and GTEx samples (Figure 1C,D) [18]. For further analysis, we designated this transcript as ARA-elncRNA1 (Androgen Receptor-Associated Enhancer lncRNA 1), and enhancer producing this transcript as CRE1 (cis-regulatory element 1)

We confirmed the interaction between AR and ARA-elncRNA1 using AR RIP-qPCR in LNCaP and 22Rv1 cell lines (Figure 1G). Additionally, we performed an RNA electrophoretic mobility shift assay (REMSA) to validate AR interaction with the predicted AR-RNA-binding motif, CUUCUCCUC (Figure 1H). The combined results from RIP-qPCR and REMSA assays demonstrate that AR in fact binds to ARA-elncRNA1.

In summary, we identified a novel lncRNA that is predominantly expressed in AR-positive tissues and cell lines and directly binds to AR. The TSS of this lncRNA is enriched with enhancer-associated features (H3K27ac, H3K4me1, PRO-Cap) and contains cis-regulatory elements from the ENCODE database. It is also annotated as an enhancer in the GeneHancer database [22]. These findings collectively establish ARA-elncRNA1 as a functional enhancer RNA associated with AR.

#### ARA-elncRNA1 is enriched in both cytoplasmic and chromatin fractions of RNA but interacts with AR only in nucleus

If ARA-elncRNA1 binds to AR, we reasoned that its expression might be AR-dependent. However, in LNCaP and 22Rv1 but not in VCaP cells, AR does not occupy the TSS of ARA-elncRNA1 (open access AR ChIP-seq data, Table S1, Figure S2A), and R1881 (synthetic testosterone) has no effect on ARA-elncRNA1 expression in LNCaP cells (Figure S2A, S2B). To investigate how ARA-elncRNA1 affects enhancer functions of the CRE1, we measured the expression of its target genes, *NUMA1* and *SHANK2*, by RT-qPCR following eRNA knockdown (KD) in LNCaP cells. Indeed, KD of eRNA significantly reduced the expression of CRE1’s target genes (Figure 2A). These findings demonstrate that expression of ARA-elncRNA1 is AR-independent and that eRNA product of CRE1 influences the expression of its target genes.

**Figure 2.**
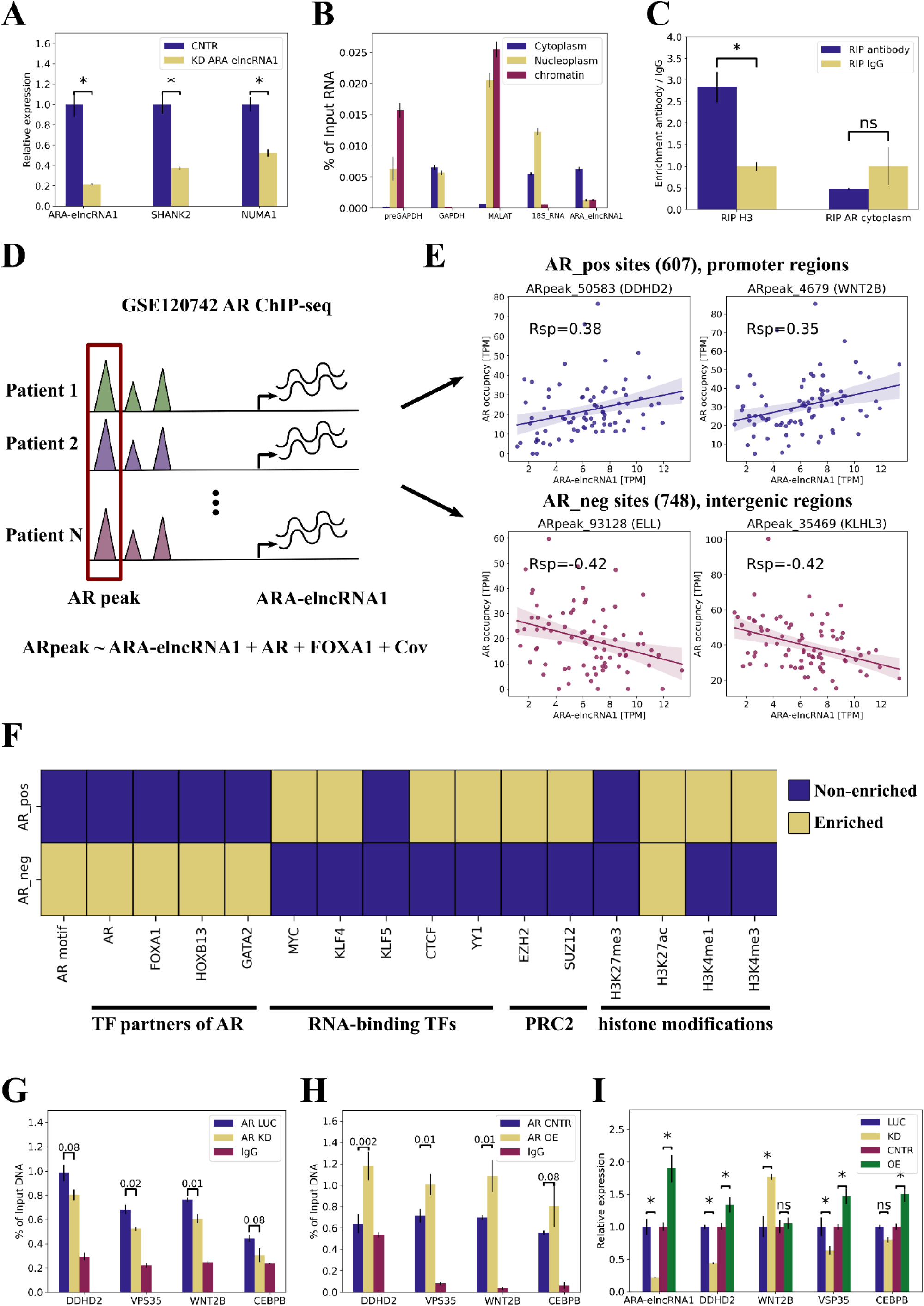
ARA-elncRNA1 is associated with AR occupancy on AR-binding sites. A. Relative normalized expression of *ARA-elncRNA1*, *NUMA1* and *SHANK2* genes following ARA-elncRNA1 KD in…cells. “*” - p-value<0.05, “ns” - non-significant. Results presented as mean ± SEM calculated from three replicates. B. The expression of *GAPDH* mRNA, *preGAPDH* (nascent transcript of GAPDH), *18S RNA*, *MALAT1* and *ARA-elncRNA1* in cytoplasm, nucleoplasm and chromatin fractions. Results presented as mean ± SEM calculated from three replicates. C. RIP results confirming the interaction between ARA-elncRNA1 and histone H3 in the nuclear fraction of LNCaP cells and absence of interaction between AR and eRNA in cytoplasmic fraction of LNCaP cells. RT-qPCR was used to examine enrichment of ARA-elncRNA1 in the anti-AR antibody fraction of RIP compared to IgG. “*” - p-value<0.05 (two-tailed unpaired Student’s t-test). Results presented as mean ± SEM calculated from three replicates. D. Schematic illustration of the computational pipeline used to find genomic regions with ARA-elncRNA1 dependent AR occupancy. E. Line plots representing correlation between AR occupancy and ARA-elncRNA expression in GSE120742 dataset for two AR_pos and AR_neg sites. The 95% confidence interval is used as an error bar. Rsp - Spearman’s rank correlation coefficient. F. Enrichment of AR_pos and AR_neg sites with ChIP-seq peaks for different chromatin associated proteins. Enrichment measured with Fisher’s exact test. Sites considered enriched if Odds Ratio > 1.5 and FDR < 0.05. PRC2 - polycomb repressive complex 2. G-H. AR occupancy of promoters of *DDHD2*, *VSP35*, *CEBPB* and *WNT2A* genes measured by qPCR-ChIP in LNCaP cells under ARA-elncRNA1 KD (G) and in 22Rv1 cells under ARA-elncRNA1 OE (H). “*” - p-value<0.05, “ns” - non-significant (two-tailed unpaired Student’s t-test). Results presented as mean ± SEM calculated from three replicates. LUC - control siRNA duplex to firefly luciferase. I. Relative normalized expression of *ARA-elncRNA1*, *DDHD2*, *VSP35*, *CEBPB* and *WNT2A* genes following ARA-elncRNA1 KD or OE in LNCaP cells. “*” - p-value<0.05, “ns” - non-significant (two-tailed unpaired Student’s t-test). Results presented as mean ± SEM calculated from three replicates.

To further explore the functions of ARA-elncRNA1, we performed cell compartment fractionation and found that this eRNA is enriched in both cytoplasmic and chromatin fractions, alongside well-known nuclear long non-coding RNA *MALAT1* and cytoplasmic *GAPDH* mRNA (Figure 2B), respectively. We confirmed the chromatin binding of ARA-elncRNA1 using RIP-qPCR with histone H3-specific antibodies (Figure 2C). Given that ARA-elncRNA1 is present in different cellular compartments, we sought to determine where AR interacts with the eRNA. Using RIP-qPCR on cytoplasmic and nuclear fractions, we found that AR interacts with ARA-elncRNA1 exclusively in the nucleus and not in the cytoplasm (Figure 1G, Figure 2C).

### ARA-elncRNA1 is associated with AR occupancy on AR-binding sites

Long non-coding RNAs (lncRNAs) can regulate transcription at various levels, ranging from direct binding to gene promoters to modulating protein stability [1, 2]. To investigate whether ARA-elncRNA1 functions as a transcriptional regulator, we analyzed open-access ChIP-seq data for AR, H3K27ac, and H3K27me3, as well as RNA-seq data from 56 patients with prostate cancer (Table S1). Using linear regression, we assessed the association between AR, H3K27ac, and H3K27me3 occupancy and ARA-elncRNA1 expression levels (Figure 2D). We found that AR occupancy at 1355 AR-binding sites (ARBS) was significantly associated with ARA-elncRNA1 expression levels, but no such associations were observed with histone marks (Figure 2E, Table S3).

It has been previously demonstrated that lncRNAs can interact with chromatin through DNA:DNA:RNA triplex formation [24–26]. To explore this possibility, we evaluated the DNA-binding potential of ARA-elncRNA1 using TriplexAligner [27]. The algorithm identified the region 1460-1484 nt of ARA-elncRNA1 as a DNA-binding domain (DBD) with high probability (Figure S2C). We further confirmed the binding of ARA-elncRNA1’s DBD to one of its predicted targets, the *CEBPB* gene promoter, using electrophoretic mobility shift assay (EMSA) (Figure S2D). Notably, the promoter of the *NUMA1* gene also contains a predicted binding site for ARA-elncRNA1, suggesting that the enhancer functions of CRE1 may also rely on ARA-elncRNA1’s ability to form RNA:DNA:DNA triplex.

The 1355 ARBS can be divided on two groups of regions: 607 regions where AR ChIP-seq signal positively correlated with ARA-elncRNA1 expression (AR_pos sites) and 748 regions where AR occupancy negatively correlated with ARA-elncRNA1 expression (AR_neg sites) (Figure 2E). These two groups exhibited distinct epigenetic and transcription factor binding profiles based on open-access ChIP-seq data (Table S1, Figure 2F). AR_pos sites were predominantly located in gene promoters (OR=14.19, p-value=1.5e-51 of the Fisher’s exact test) and enriched with H3K27ac and H3K4me3 signals (Figure 2E, F). In contrast, AR_neg sites were primarily located in intergenic regions (OR=1.34, p-value=0.03 of the Fisher’s exact test) and avoided promoters (OR=0.34, p-value=2.8e-8 of the Fisher’s exact test). AR_neg sites were enriched for ChIP-seq peaks of well-known PCa regulators, including AR, FOXA1, HOXD13, and GATA2 (Figure 2F, F) [28]. Conversely, AR_pos sites were enriched for ChIP-seq signals of proteins with known RNA-binding activity, such as YY1, CTCF, EZH2, SUZ12, MYC, and KLF4 (Figure 2F) [6, 29–32]. Interestingly, AR_pos sites lacked AR-binding motifs despite being derived from AR ChIP-seq data (Figure 2F). We also analyzed open-access RNA-seq data from DHT(dihydrotestosterone)-treated LNCaP, 22Rv1, and VCaP cells, as well as from LNCaP and VCaP cells with *AR* knockdown, and PCa cells with KD of *FOXA1*, *HOXD13*, *GATA2*, *YY1*, and *EZH2* (Figure S2E, Table S1). We found that ARBS were generally located in genes whose expression changed under these conditions (Figure S2E, Table S3). Genes located near AR_pos sites (referred to as AR_pos genes) were significantly enriched in Reactome pathways such as “WNT Ligand Biogenesis and Trafficking,” “RHO GTPase Cycle,” “Cellular Responses to Stimuli,” and “RNA Polymerase II Transcription” (Figure S2F). Genes involved in the WNT and RHO GTPase Cycle pathways play critical roles in processes such as cell migration, adhesion, division, establishment of cellular polarity, and intracellular transport. This suggests that ARA-elncRNA1 may potentially regulate these aspects of cell behavior.

To investigate the influence of ARA-elncRNA1 on AR occupancy of ARBS, we selected four AR_pos sites located in the promoters of *DDHD2*, *VSP35*, *WNT2B*, and *CEBPB* genes, as well as two AR_neg sites located in *ELL* and *KLHL3* genes, based on the correlation between AR ChIP-seq signal and ARA-elncRNA1 expression (Table S3). ARA-elncRNA1 knockdown resulted in a decrease in AR occupancy at the promoters of *WNT2B* and *VSP35* genes, as measured by ChIP-qPCR (Figure 2G). Conversely, overexpression of a fragment of ARA-elncRNA1 containing ARE and DBD (Figure S2G) in 22Rv1 cells increased AR occupancy at the promoters of *DDHD2*, *WNT2B*, and *VSP35* genes (Figure 2H). Notably, AR occupancy at the *DDHD2* promoter in 22Rv1 cells was initially indistinguishable from the IgG signal, but overexpression of ARA-elncRNA1 doubled the AR ChIP signal (Figure 2H).

We also tested the effect of knockdown and overexpression of eRNA on the expression of these genes (Figure 2I, Figure S2H). In LNCaP cells, *DDHD2* and *VSP35* expression levels changed in accordance with ARA-elncRNA1 levels. *WNT2B* expression increased under eRNA KD but showed no change under OE. *CEBPB* expression increased under OE but remained unchanged under KD. Interestingly, ARA-elncRNA1 knockdown in 22Rv1 cells had no effect on the expression of these genes, while overexpression of ARA-elncRNA1 increased the expression of only the *DDHD2* gene (Figure 2I, Figure S2H). These findings indicate that ARA-elncRNA1 not only influences AR occupancy at the promoters of AR_pos genes but also regulates their expression.

We also tested two AR_neg sites and found that ARA-elncRNA1 knockdown led to increased AR occupancy at an AR_neg region in the *KLHL3* gene, whereas overexpression of ARA-elncRNA1 had little to no effect on AR occupancy of this region (Figure S2I, S2J).

#### ARA-elncRNA1 regulates expression of AR_pos genes through AR:ARA-elncRNA1:YY1 axis

As we have shown, ARA-elncRNA1 influences AR occupancy at certain AR-binding sites. However, the molecular mechanism underlying this influence remains unclear. We know that AR_pos sites are enriched with RNA-binding transcription factors such as YY1, CTCF, and PRC2 proteins (Figure 2F, S3A) but lack AR binding motifs (Figure 2F). We hypothesize that AR interacts with AR_pos sites not through direct binding to DNA motif but via ARA-elncRNA1, which may bind to chromatin or RNA-binding proteins.

To test this hypothesis, we performed native RIP experiments in 22Rv1 cells using antibodies against YY1, EZH2, and SUZ12. Our results confirmed an interaction between ARA-elncRNA1 and YY1 (Figure 3A) but not with EZH2 or SUZ12 (Figure S3B). Additionally, we validated the interaction between YY1 and eRNA in LNCaP cells (Figure 3A) and the absence of interaction with EZH2 using an EZH2 RIP-seq datasets (Table S1, Figure S3C). Furthermore, we confirmed the interaction of YY1 with four selected AR_pos regions through ChIP-qPCR in both LNCaP and 22Rv1 cells (Figure 3B). We also demonstrated the interaction of ARA-elncRNA1 with chromatin regions of four AR_pos loci using Chromatin Isolation by RNA Purification (ChIRP, Figure 3C).

**Figure 3.**
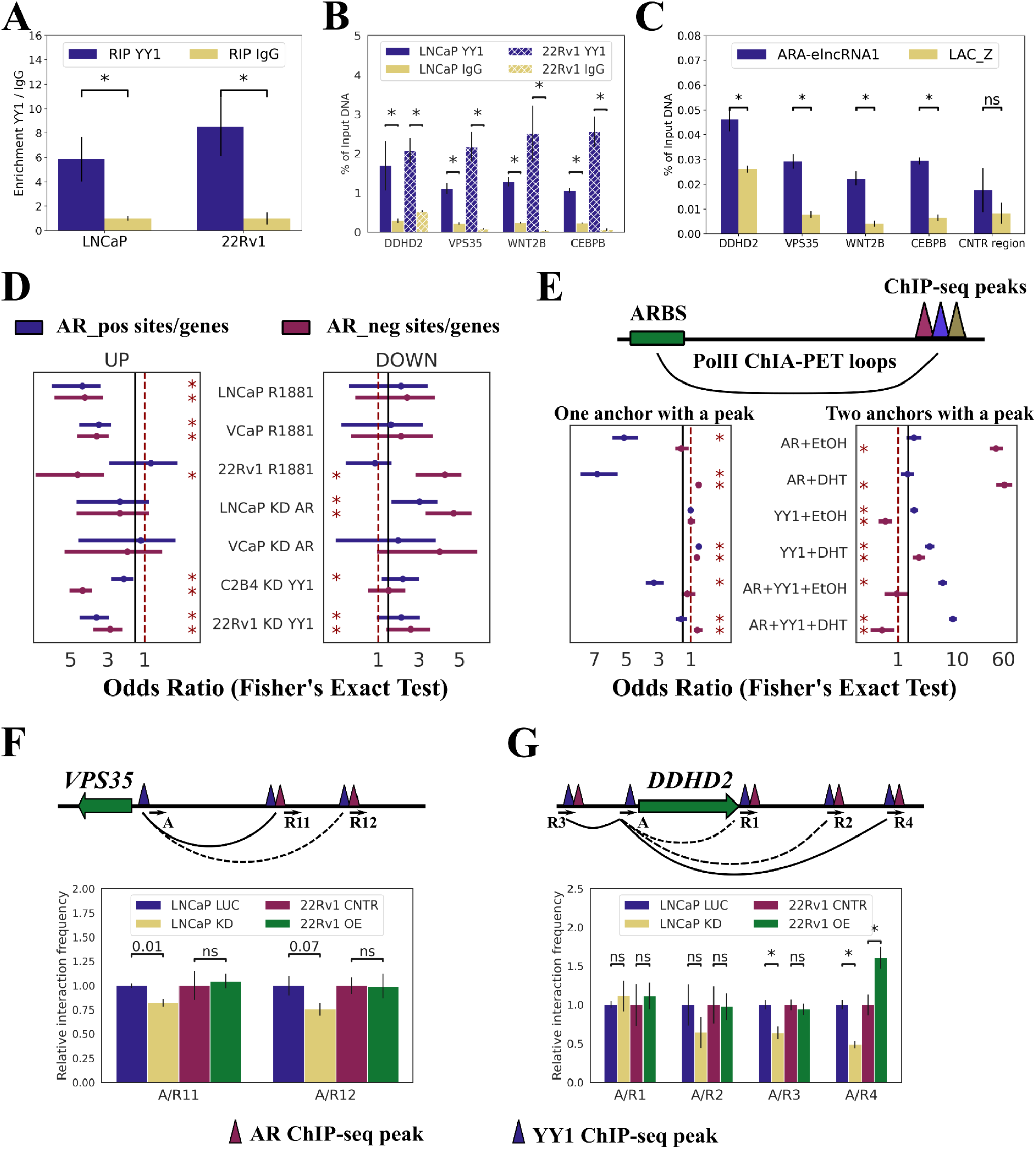
ARA-elncRNA1 regulates AR_pos gene expression through the AR:ARA-elncRNA1:YY1 axis. A. RIP results confirming the interaction between ARA-elncRNA1 and YY1 in the nuclear fraction of LNCaP and 22Rv1 cells. RT-qPCR was used to assess the enrichment of ARA-elncRNA1 in the anti-AR antibody fraction compared to IgG. “*” - p-value < 0.05 (two-tailed unpaired Student’s t-test). Results are presented as mean ± SEM from three replicates. B. YY1 occupancy at the promoters of *DDHD2*, *VSP35*, *CEBPB*, and *WNT2A* genes was measured by ChIP-qPCR in LNCaP and 22Rv1 cells. “*” - p-value < 0.05 (two-tailed unpaired Student’s t-test). Results are presented as mean ± SEM from three replicates. C. ARA-elncRNA1 interactions with the promoters of *DDHD2*, *VSP35*, *CEBPB*, and *WNT2A* genes were measured by ChIRP-qPCR in LNCaP cells. Antisense oligos tiling ARA-elncRNA1 and LacZ (negative control) were used. As a control region, the AR binding site on the 3’ end of *DDHD2* gene was selected (ARpeak_50586, Table S3). “*” - p-value < 0.05 (two-tailed unpaired Student’s t-test). Results are presented as mean ± SEM from three replicates. D. Enrichment of AR_pos and AR_neg genes with up-and down-regulated genes following R1881 treatment, AR KD, and YY1 KD in corresponding cell lines. Odds ratios with 95% confidence intervals are shown. “*” - p-value < 0.05 (Fisher’s exact test). E. Schematic illustration of the ChIA-PET loops data analysis pipeline used to identify genomic regions with ARA-elncRNA1-dependent AR occupancy. Odds ratios with 95% confidence intervals are shown. “*”-p-value < 0.05 (Fisher’s exact test). F-G. 3C-qPCR assays were performed using crosslinked, MboI-digested chromatin from LNCaP cells following ARA-elncRNA1 KD or 22Rv1 cells after OE of ARA-elncRNA1, treated with 10 nM R1881 for 24 h. The black arrow indicates the forward primer near the MboI restriction site, and’A’ denotes the anchor region. Bar plots show normalized E-P interactions for *VSP35* (F) and *DDHD2* (G) genes. Solid black loops indicate statistically significant changes in E-P interactions, while dashed black loops indicate non-significant changes. “*” - p-value < 0.05 (two-tailed unpaired Student’s t-test), “ns” - non-significant. Results are presented as mean ± SEM from three replicates. LUC - control siRNA duplex to firefly luciferase.

These findings, coupled with our data showing ARA-elncRNA1’s influence on AR interaction with AR_pos sites, suggest that this eRNA facilitates AR recruitment to AR_pos sites occupied by YY1. As previously demonstrated, changes in ARA-elncRNA1 abundance lead to altered expression of AR_pos genes (Figure 2I, S2H). Additionally, analysis of open-access RNA-seq datasets from YY1 and AR knockdowns or R1881 treatment in LNCaP, VCaP, and 22Rv1 cells revealed significant changes in the expression of AR_pos genes (Figure 3D, S2E). Together, these results support the hypothesis that the AR:ARA-elncRNA1:YY1 complex regulates the expression of AR_pos genes.

It is well-established that YY1 facilitates chromatin looping between enhancers and promoters (E-P loops) [33]. Using open-access PolII ChIA-PET data from LNCaP and VCaP cells (Table S1), we confirmed that AR_pos sites are enriched with loops where only one anchor intersects an AR ChIP-seq peak, but not both anchors (Figure 3E, OR=6.8, p-value=4.2e-124). In contrast, AR_neg sites are significantly enriched with loops framed by AR peaks on both anchors (Figure 3E, OR=61.7, p-value=3.5e-215). AR_pos sites are depleted in loops anchored on a YY1 ChIP peak on one side (Figure 3E, OR=0.5, p-value=1.2e-16) but enriched in loops anchored on YY1 peaks on both sides (Figure 3E, OR=3.4, p-value=7.1e-50). Interestingly, loops simultaneously anchored on AR and YY1 ChIP peaks are more likely to contain AR_pos sites than loops with only one YY1 or AR peak (Figure 3E, OR=8.4 vs. OR=3.4 and OR=6.8, p-value<1e-40, Fisher’s exact test). This suggests that ARA-elncRNA1 may influence chromatin looping.

To explore this further, we selected two genes with AR_pos sites located in their promoters: *DDHD2* and *VPS35*. For *DDHD2*, we identified four nearby or connected regions (anchors) through ChIA-PET loops, which were occupied by both AR and YY1. For *VPS35*, two regions meeting the same criteria were selected. Using chromatin conformation capture (3C) analysis [34], we demonstrated that ARA-elncRNA1 knockdown in LNCaP cells reduced interactions between the *VPS35* promoter and region R11 (Figure 3F), as well as between the *DDHD2* promoter and regions R3 and R4 (Figure 3G). Conversely, overexpression of ARA-elncRNA1 in 22Rv1 cells increased interactions between the *DDHD2* promoter and region R4 (Figure 3G). These findings support the hypothesis that ARA-elncRNA1 facilitates chromatin looping between YY1-occupied promoters and AR-occupied distant regions by interacting with both AR and YY1.

Together, these results indicate that ARA-elncRNA1 influences chromatin looping by assembling an AR:ARA-elncRNA1:YY1 complex on E-P loops.

#### ARA-elncRNA1 modulates AR abundance at mRNA and protein levels

Another potential mechanism by which ARA-elncRNA1 regulates AR occupancy at ARBS is through the modulation of AR expression. KD of eRNA decreased the expression of both *AR* and *KLK3* genes, regardless of R1881 treatment (Figure 4A). Conversely, OE of eRNA increased *AR* expression independently of R1881 (Figure S3D). Interestingly, overexpression of ARA-elncRNA1 had a similar effect on *KLK3* expression as R1881 treatment (Figure S3D). Additionally, when comparing the effect of R1881 on LNCaP cells with ARA-elncRNA1 KD, it appears that AR activation does not compensate for the KD effect but rather enhances it (Figure 3SE), especially in the case of *CEBPB* gene.

**Figure 4.**
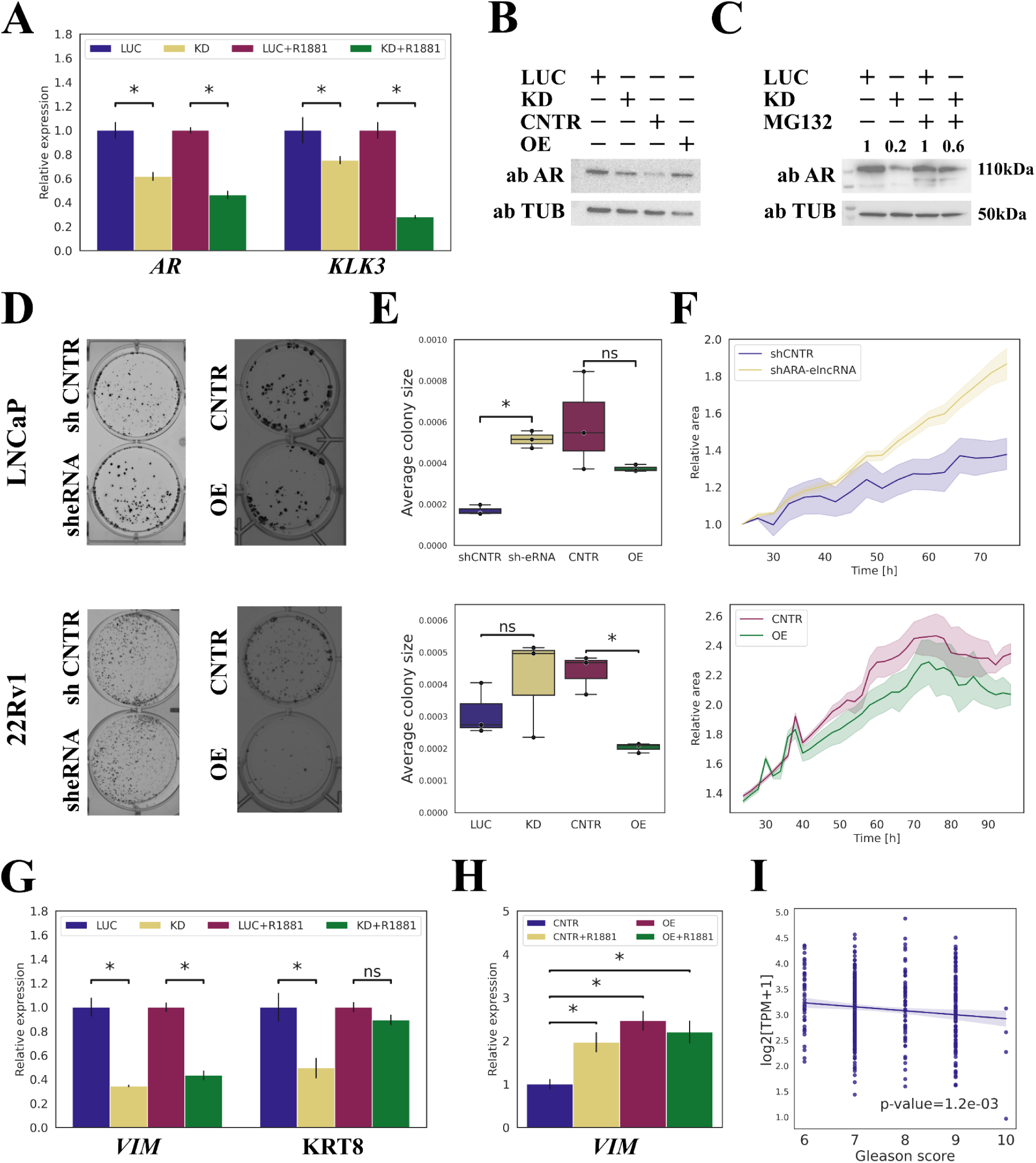
ARA-elncRNA1 modulates AR expression and influences clonogenicity and proliferation in PCa cell lines. A. Relative normalized expression of *AR* and *KLK3* genes following ARA-elncRNA1 KD and R1881 treatment. “*” - p-value<0.05, “ns” - non-significant (two-tailed unpaired Student’s t-test). Results presented as mean ± SEM calculated from three replicates. B. Western blot analysis of LNCaP following KD/OE of ARA-elncRNA1 cell lysates using anti-AR or anti-TUB antibodies. Numbers indicate relative abundance over control (LUC) of the protein. LUC - control siRNA duplex to firefly luciferase. C. Western blot analysis of LNCaP under KD of ARA-elncRNA1 treated or untreated with MG132 (12.5 uM) for 4.5 h cell lysates using anti-AR or anti-TUB antibodies. Numbers indicate relative abundance over control (LUC) of the protein. LUC - control siRNA duplex to firefly luciferase. D-E. Colony formation assay of LNCaP or 22Rv1 cells under KD/OE of ARA-elncRNA1. Representative images of colonies in wells are shown in panel D. Box-plots representing the average size of a colony are shown in panel E. Each experiment performed in three biological replicates. “*” - p-value<0.05, “ns” - non-significant (two-tailed unpaired Student’s t-test). F. KD and 22Rv1-mCherry cells under OE of ARA-elncRNA1. Line plot shows the relative area occupied by cells, measured using the Celena X High Content Imaging System over 48-100 hours. Images were acquired every 2-3 hours. The 95% confidence interval is used as an error bar. Each experiment performed in three biological replicates. G. Relative normalized expression of *VIM* and *KRT8* genes under ARA-elncRNA1 KD and R1881 treatment. “*” - p-value<0.05, “ns” - non-significant (two-tailed unpaired Student’s t-test). Results presented as mean ± SEM calculated from three replicates. LUC - control siRNA duplex to firefly luciferase. H. Relative normalized expression of *VIM* gene under ARA-elncRNA1 OE and R1881 treatment. “*” - p-value<0.05 (two-tailed unpaired Student’s t-test). Results presented as mean ± SEM calculated from three replicates. LUC - control siRNA duplex to firefly luciferase. I. Line plot showing a negative correlation between ARA-elncRNA1 expression levels and Gleason score, based on TCGA PRAD data analysis (Ordinary Least Squares regression, p-value<0.05).

Western blot (WB) analysis of LNCaP and 22Rv1 cells following ARA-elncRNA1 KD revealed a decrease in AR protein levels (Figure 4B, Figure S3F). In contrast, overexpression of eRNA led to AR stabilization (Figure 4B). It is known that some lncRNAs can protect proteins including AR from proteasomal degradation [35, 36]. The addition of the proteasomal inhibitor MG132 abolished the effect of ARA-elncRNA1 knockdown on AR protein levels (Figure 4C, Figure S3F). These findings suggest that ARA-elncRNA1 protects AR from proteasomal degradation.

These results align with the observed association between ARA-elncRNA1 expression and AR occupancy at AR_pos sites. However, how the decrease in AR protein levels results in increased AR occupancy at AR_neg sites remains unclear. This phenomenon may be due to competition between AR binding to classical DNA motifs and its interaction with chromatin through ARA-elncRNA1. Although the reduction of eRNA leads to a decrease in AR levels, the remaining AR molecules may interact more strongly with classical AR binding sites.

### ARA-elncRNA1 exerts antitumorigenic effects

To investigate how ARA-elncRNA1 affects PCa cell survival, we performed two experiments: a clonogenic assay and monitoring of cell confluency over time. We used LNCaP and 22Rv1 cell lines expressing mCherry as a fluorescent marker, with stable knockdown of ARA-elncRNA1 achieved through shRNA interference.

Knockdown of ARA-elncRNA1 by shRNA increased the average colony area compared to the control but decreased the clonogenic survival of LNCaP cells (Figure 4D, E, upper panel). In contrast, no significant effect was observed in 22Rv1 cells (Figure 4D, E, bottom panel). Overexpression of ARA-elncRNA1 in both LNCaP (Figure 4D, E, upper panel) and 22Rv1 (Figure 4D, E, bottom panel) cell lines reduced the average colony size.

Additionally, LNCaP cells with stable KD of ARA-elncRNA1 showed significantly increased confluency compared to control cells (Figure 4F, upper panel). However, the effect of ARA-elncRNA1 KD using siRNA differed from that of shRNA-mediated KD in LNCaP cells. Under siRNA KD, the average colony size for LNCaP cells was smaller than in control cells (Figure S3G-H). Cell confluency was higher in siRNA KD cells than in control cells in a period between 30 and 40 hours, after which confluency became similar in both conditions (Figure S3I). This transient effect is likely due to the limited activity of siRNA oligonucleotides in cells.

In 22Rv1 cells, siRNA-mediated KD of ARA-elncRNA1 had no effect on average colony size (Figure S3J-K) or confluency (Figure S3L). However, overexpression of ARA-elncRNA1 in 22Rv1 cells slightly but significantly decreased cell confluency (Figure 4F, bottom panel).

We hypothesized that ARA-elncRNA1 might influence the mesenchymal state of cells and therefore tested the expression of mesenchymal markers: *KRT8* and the transcription factor *VIM*. Knockdown of ARA-elncRNA1 decreased the expression levels of these genes in…cells, while overexpression of ARA-elncRNA1 increased their expression (Figure 4G, H). However, the addition of R1881 to the cells abolished the effects of ARA-elncRNA1 knockdown (Figure 4G, H). This suggests that ARA-elncRNA1 does not directly regulate mesenchymal genes but may influence them through its interaction with AR, as we have previously demonstrated. Analysis of the TCGA prostate cancer (PCa) patient cohort data revealed that high expression levels of ARA-elncRNA1 are associated with a lower Gleason score but not with survival outcomes (Figure 4I).

These results are consistent with the clonogenic, confluency assays and TCGA data, indicating that ARA-elncRNA1 is required for maintaining cellular properties associated with a less aggressive phenotype in prostate cancer cells.

## Discussion

Here, we investigated whether there are any long non-coding RNAs expressed from enhancer regions that interact with the androgen receptor in a sequence-specific manner and play a significant role in prostate cancer development. To address this question, we analyzed open-access RNA-seq and AR-RIP-seq data to identify novel intergenic transcripts that potentially bind to AR. Our analysis revealed that only one transcript expressed from an enhancer region met stringent filtering criteria: the transcript’s transcription start site (TSS) overlapped with enhancer features such as H3K27ac, H3K4me1, and GRO-cap peaks, and it was expressed in AR-positive prostate cancer (PCa) cell lines and TCGA PRAD sample collection. We named this transcript ARA-elncRNA1.

In a pioneering study of AR-binding RNAs, researchers identified several hundred unannotated lncRNAs that might interact with AR [14]. However, that study did not distinguish between lncRNAs and enhancer RNAs (eRNAs), nor did it apply stringent filtration criteria such as RNA motif analysis. Consequently, in our study, we identified only one strong candidate eRNA. We subsequently confirmed that this eRNA interacts with AR in a sequence-specific manner within the cell nucleus but not in the cytoplasm, and it contains DNA:DNA:RNA triplex forming region (DNA-binding region). Previous studies have shown that lncRNAs can target AR to gene promoters and activate gene expression [12, 13]. For example, the lncRNA SLNCR1 forms a complex with AR and transcription factor Brn3a to bind to the *MMP9* promoter [12]. This led us to ask: Could ARA-elncRNA1 recruit AR to gene promoters?

Analysis of open-access data allowed us to predict 1355 regions where AR occupancy levels correlate with ARA-elncRNA1 expression. These regions were divided into two groups: AR_pos, which showed a positive correlation with ARA-elncRNA1 expression, and AR_neg, which showed a negative correlation. These two groups exhibited distinct histone modification and transcription factor binding profiles. AR_pos regions lacked AR DNA-binding motifs, were located in promoters, and were enriched with ChIP-seq peaks of RNA-binding TFs and proteins such as YY1 and EZH2 [29, 30, 37]. YY1 is a transcription factor known to mediate enhancer-promoter interactions and has nonspecific RNA-binding capabilities [33]. Previous studies have shown that lncRNAs can shuttle other proteins to YY1 binding sites [29]. In contrast to AR_pos regions, AR_neg regions contained AR DNA-binding motifs, were located in intergenic regions, and were enriched with ChIP-seq peaks of PCa-associated TFs, such as FOXA1 and HOXB13. We confirmed that knockdown and OE of ARA-elncRNA1 increased and decreased AR occupancy at AR_pos sites and AR_neg, respectively, as measured by AR ChIP-qPCR. Conversely, KD of ARA-elncRNA1 increased AR binding to AR_neg site.

The next question was: What is the possible molecular mechanism by which ARA-elncRNA1 influences AR binding to AR_pos and AR_neg regions? The enrichment of AR_pos sites with RNA-binding proteins suggested that ARA-elncRNA1 might shuttle AR to one of these proteins at gene promoters. Indeed, RIP-qPCR, ChIP-qPCR, and ChIRP experiments confirmed that ARA-elncRNA1 interacts not only with AR but also with YY1. Moreover, YY1 binds to AR_pos regions, as does ARA-elncRNA1 itself. We also demonstrated that KD and OE of ARA-elncRNA1 influences looping between enhancers of AR_pos genes and distal enhancers co-occupied by AR and YY1, and alter the expression of AR_pos genes. Collectively, these findings indicate that ARA-elncRNA1 regulates the expression of several AR_pos genes by targeting AR to YY1 and stabilizing enhancer-promoter (E-P) loops.

Interestingly, ARA-elncRNA1 not only recruits AR to promoters but also regulates AR expression at both the RNA and protein levels. We showed that KD of ARA-elncRNA1 reduces AR abundance, but this effect iis abolished by proteasome inhibition with MG132. This suggests that ARA-elncRNA1 protects AR from proteasomal degradation. Some lncRNAs are known to protect proteins from degradation by blocking their interaction with E3 ubiquitin ligases [35, 36]. This protective effect of ARA-elncRNA1 on AR protein levels may explain the increased AR occupancy at AR_pos sites. However, we did not observe compensation of the ARA-elncRNA1 KD effect on AR_pos gene expression upon testosterone treatment in LNCaP cells. Testosterone induces AR dimerization and translocation to the nucleus at high levels [11], but in the absence of ARA-elncRNA1, this has no effect on AR_pos gene expression. Despite the decrease in AR protein levels under eRNA KD, AR binding to AR_neg sites increases. We may explain this discrepancy using a competition model: although AR abundance is lower under ARA-elncRNA1 KD, the absence of ARA-elncRNA1 prevents AR from being shuttled away from sites with DNA-binding motifs to AR_pos regions. Another possible scenario is that ARA-elncRNA1 forms liquid-liquid phase separation condensates that sequester AR through its AR motif, preventing interaction with AR_neg sites and possibly proteasomes. Although we need more data to confirm any of these theories.

Interestingly, the two AR+ cell lines we tested exhibited distinct outcomes in our KD or OE experiments. LNCaP cells responded more strongly to KD treatment—using either shRNA or siRNA—than 22Rv1 cells, as evidenced by changes in AR_pos gene expression levels, AR occupancy, E-P looping, and cell proliferation. This difference may be attributed to the varying expression levels of endogenous ARA-elncRNA1: qPCR measurements revealed that 22Rv1 cells express 8 times less ARA-elncRNA1 than LNCaP cells. Additionally, in OE experiments, 22Rv1 cells showed some effects, though less pronounced. Another potential factor is the difference in AR protein isoforms between the two cell lines. While LNCaP cells express full-length AR, western blot analysis of 22Rv1 cells revealed two bands: the higher band corresponding to full-length AR and the lower band corresponding to the ARv7 isoform, which lacks the ligand-binding domain (LBD). However, how the absence of the LBD might affect transcriptional activity of the ARA-elncRNA1:AR complex remains unclear.

In summary, we have identified a novel enhancer-associated long non-coding RNA (eRNA) that binds to the androgen receptor in a sequence-specific manner, shuttles it to YY1-occupied promoters, and stabilizes E-P loops (Figure 5). This eRNA influences AR protein stability and directly regulates AR occupancy at several genomic regions. In LNCaP cells, KD of this eRNA significantly increases cell proliferation. To our knowledge, this is the first report of an eRNA that trans-regulates gene expression through interactions with AR and YY1. These findings may provide new insights into the complexity of androgen-dependent transcriptional networks and the mechanisms underlying cancer development.

**Figure 5.**
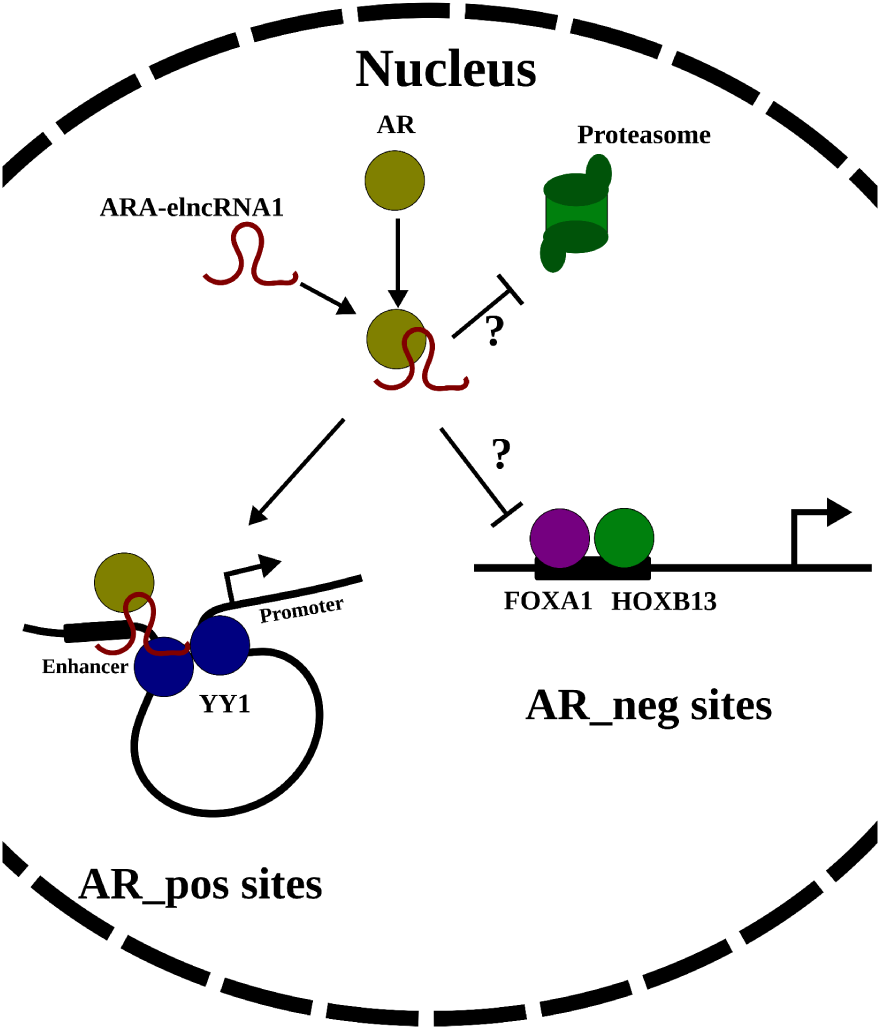
Model of ARA-elncRNA1 function in AR-dependent gene expression ARA-elncRNA1 binds to AR in a sequence-specific manner and protects it from proteasomal degradation. The AR:ARA-elncRNA1 complex does not bind to classical AR enhancers but instead shuttles to YY1-mediated enhancer-promoter loops, activating the expression of AR-dependent genes.

## Materials and Methods

### RNA-seq data analysis

For *de novo* transcript assembly, all datasets from Table S1 were merged into a single FASTQ. The resulting FASTQ files were mapped to the human genome (hg38) using the STAR 2.7.10a program with standard parameters. Samtools 1.15.1 was used for BAM file manipulations. The Scallop2 algorithm was employed for *de novo* assembly with the following command: “scallop-i $input_bam-o $output_file--verbose 0--min_single_exon_coverage 10”. A FASTA file containing the sequences of *de novo* transcripts was generated using the gffread v0.12.7 function, with the GFF output file from Scallop and the human reference genome as inputs.

For differential expression (DE) analysis, FASTQ files were trimmed using Trimmomatic 0.39 and pseudoaligned to the reference transcriptome (GENCODE v37) with kallisto 0.48.0. Gene expression tables were generated using the tximport tool. DE analysis was performed using DESeq2. Genes with |log2FC| > 0.5 and FDR < 0.05 were considered differentially expressed. Here, FC (fold change) represents the ratio of expression in the experimental condition to the control condition, and FDR (false discovery rate) was calculated using the Benjamini-Hochberg procedure for multiple testing correction of p-values.

For *de novo* transcript coverage calculation, kallisto 0.48.0 and tximport were used, with the sequences of *de novo* transcripts used as the reference.

For the parallelization of calculations, GNU Parallel was utilized [38].

### Gro-seq and Gro-Cap data analysis

Reads were trimmed using Trimmomatic 0.39. Reads originating from ribosomal genes were filtered out using the BBDuk program. The remaining reads were mapped to the human reference genome (hg38) using bwa mem 0.7.17-r1188. BigWig files representing coverage were generated using deepTools2 bamCoverage. Transcription start site (TSS) regions were identified using the pints program [39].

### ChIP-seq data analysis

Reads were trimmed using Trimmomatic 0.39. The remaining reads were mapped to the reference genome (hg38) using bwa mem 0.7.17-r1188. ChIP-seq peaks were called using the MACS3 program with standard parameters. BigWig files containing coverage data were generated using deepTools2 bamCoverage with a sliding window of 50 bp. Bed-files for LNCaP ChIP-seq MYC and CTCF samples were downloaded from cistrom.db [40].

### ChIA-PET data analysis

Bedpe files for LNCaP (GSM3423998) and VCaP (GSM3423997) were downloaded from GEO (Table S1). The genomic coordinates of the loops were converted from the hg19 to the hg38 genome version using the UCSC liftOver tool [41]. The two files were merged using the hicMergeLoops function from the HiCExplorer Python package [42] with the following parameters:--inputFiles VCaP.hg38.bedpe LNCaP.hg38.bedpe--outFileName merged_VCaP_LNCaP.bedpe-r 2500.

### DNA motif analysis

AR DNA motif analysis was conducted using the FIMO program with the HOCOMOCO v11 motif database.

### Association analysis between ChIP-seq signal and ARA-elncRNA1 expression

Dataset GSE120742 contains ChIP-seq data for AR, H3K27ac, and H3K27me3, as well as RNA-seq data from prostate cancer (PCa) patients. ChIP-seq datasets were analyzed as previously described, with minor modifications: for MACS3 peak calling, a q-value threshold of 0.1 was used. The resulting peak files from AR, H3K27ac, and H3K27me3 ChIP-seq experiments were merged into a single file using BEDTools merge (v2.31.1). Coverage of each region in the BED file was calculated using BEDTools coverage for each ChIP-seq BAM file. The coverage of each region was normalized by library size and region length.

RNA-seq data were pseudoaligned to either the reference transcriptome (gencode.v37) or the *de novo* transcriptome (LNCaP_scallop). Gene coverage was normalized using TPM (Transcripts Per Kilobase Million).

For the final analysis, only data from patients with all four types of experiments available were used (56 patients). ChIP-seq and RNA-seq data matrices were min-max normalized. The RNA-seq data matrix was subjected to principal component analysis (PCA), and the first four principal components were included as covariates in the regression models.

To study the association between AR occupancy and ARA-elncRNA1 expression, the following three linear regression models were used:

1. AR_peak_i_ = const + E(ARA-elncRNA1) + E(AR) + E(FOXA1) + Cov;
2. H3K27ac_peak_i_ = const + E(ARA-elncRNA1) + E(AR) + E(FOXA1) + Cov;
3. H3K27me3_peak_i_ = const + E(ARA-elncRNA1) + E(EZH2) + E(SUZ12) + Cov; where:

const: constant term; AR_peak_i_ / H3K27ac_peak_i_ / H3K27me3_peak_i_: normalized ChIP-seq signal vector for the i-th region;

E(Gene Name): normalized expression vector of the specified gene;

Cov: PC1–PC4 (the first four principal components from PCA of the gencode.v37 RNA-seq matrix).

Expression levels of AR, FOXA1, EZH2, and SUZ12 were included as covariates to minimize false-positive associations. Linear regression was performed using the statsmodels.OLS (v0.13.2) Python library. PCA and data normalization were conducted using modules from the sklearn (v1.0.2) Python library.

### Enrichment analysis

To test for non-random intersections between lists of genes or regions, Fisher’s exact test was used. Significance criteria were defined as an odds ratio (OR) > 1.5 and a p-value (or FDR-adjusted p-value if multiple tests were performed) < 0.05.

For enrichment analysis of Reactome pathways, the enricher function from the clusterProfiler R library was used, with reference gene lists from MSigDB.

### Cell lines and cell culture

LNCaP (American Type Culture Collection, CRL-1740), 22Rv1 (American Type Culture Collection, CRL-2505) and Phoenix-GP (ATCC-CRL-3215) cells were used in experiments. LNCaP and 22Rv1 were cultured in RPMI medium (PanEco, Russia) supplied with 10% (v/v) FBS (Gibco Fetal Bovine Serum, United States), GlutaMAX Supplement (Thermo Fisher Scientific, USA) and gentamicin (10 ug/ml, PanEco, Russia). Phoenix-GP cells were cultured in DMEM (PanEco, Russia) + 10% FBS + GlutaMAX Supplement + gentamicin (10 ug/ml). All cell lines were incubated at 37 °C and 5% CO_2_ in a CO_2_ incubator. For some experiments LNCaP and 22Rv1 cells were treated with 10 uM of R1881 (synthetic testosterone, R0908, Sigma-Aldrich, USA) for 24h. For proteasome inhibition cells were treated with MG132 (Thermo Fisher Scientific USA) 12.5uM for 4.5 h.

### Plasmids, lentivirus production and cell transduction

Construction of ARA-elncRNA1knockdown vector The pLKO.1_puro plasmid (Addgene #421039) was used to knockdown ARA-elncRNA1 via siRNA interference, following the Addgene protocol for shRNA cloning with minor modifications (https://www.addgene.org/protocols/plko/). Briefly, single-stranded DNA (ssDNA) templates were synthesized by Evrogen (Russia) and annealed in 1× NEB2 buffer (New England Biolabs, UK) to form double-stranded DNA (dsDNA). The vector was digested with AgeI and EcoRI restriction enzymes (Fermentas), and the dsDNA template was ligated into the linearized vector using T4 DNA ligase (Evrogen, Russia). The ligation mixture was transformed into *E. coli* strain XL1-Blue (Evrogen, Russia). The pLKO.1-puro Non-Mammalian shRNA plasmid (Merck, SHC002) was used as a control.

ssDNA Templates:

gene56338sh1F:5’-CCGGAACATGGCAGTACAAATATCTTTCTCGAGAAAGATATTTGTACTGCCATGTTTTTTTCTAGAG-3’,

gene56338sh1R:5’-AATTCTCTAGAAAAAAACATGGCAGTACAAATATCTTTCTCGAGAAAGATATTTGTACTGCCATGTT-3’.

Construction of ARA-elncRNA1 expression vector

An ARA-elncRNA1 fragment containing the AR-binding site and predicted DNA-binding region was amplified using the following primer pair:

Forward Primer: 5’-ttgctagcAGCATTCAAATCTGCAGACCGA-3’, Reverse Primer: 5’-tttctagaTGGGGGAGGAGGAAACATAGAG-3’.

LNCaP cDNA was used as the template. The PCR product, containing NheI/XbaI restriction sites (highlighted in lowercase in the primer sequences), was ligated into the linearized pcDNA3.1/hygro vector (Invitrogen, V87020) to generate the ARA-elncRNA1 overexpression plasmid. The plasmid was verified by sequencing. The empty pcDNA3.1/hygro vector was used as a control in overexpression experiments.

Producing Lentiviral Particles

For pLKO.1 backbone (Addgene #421039):

Lentiviral particles were produced using the psPAX2 (Addgene #12260) and pMD2.G (Addgene #12259) packaging plasmids. Briefly, Phoenix-GP cells were seeded in a T25 flask to reach 50–60% confluency by the next morning. The cells were transfected with a mixture of 4 µg of the target plasmid, 3 µg of psPAX2, and 1 µg of pMD2.G using Turbofect (Thermo Fisher Scientific, R0531) according to the manufacturer’s protocol. The medium was replaced the following day, and after 3 days, the virus-containing medium was collected, filtered through a 0.45 µm filter, aliquoted into 500 µL portions, and stored at –70 °C.

For LeGO-iC backbone (Addgene #27362):

Lentiviral particles were produced using auxiliary plasmids containing the Rev (15.3% by mass of total plasmid DNA), RRE (29.8% by mass of total plasmid DNA), and VSV-G (5.6% by mass of total plasmid DNA) genes. Phoenix-GP cells were transfected with a mixture of all four plasmids. The subsequent steps were identical to the protocol described above for the pLKO.1 backbone.

Cell lines transduction

Cells were seeded in a 6-well plate at a density of 100,000 cells per well. The following day, one aliquot of thawed virus was added to the medium (in a 1:1 ratio) in each well, along with polybrene (10 ug/mL). After 24 hours, the medium was replaced with a fresh full medium without virus.

For LNCaP cells transduced with viruses on the pLKO.1 backbone, puromycin was added to the medium at a concentration of 2.5 ug/mL for 4 days. For LNCaP and 22Rv1 cells transduced with LeGO-iC viruses, cells were sorted based on mCherry fluorescence signal (BD FACSMelody™ Cell Sorter, USA).

### Knockdown and overexpression experiments

ARA-elncRNA1 knockdown was performed using RNA interference. RNA oligonucleotides (5’:GUUGAAGAGGAGAGAAUUAUU, 3’:UUCAACUUCUCCUCUCUUAAU) were designed using a siRNA selection program from the Whitehead Institute (https://sirna.wi.mit.edu/) and synthesized by Syntol (Russia). Oligos were annealed in siRNA resuspension buffer (1x: 60 mM KCl, 6 mM HEPES-KOH, pH 7.5, 0.2 mM MgCl_2_). siRNA was transfected into LNCaP or 22Rv1 cells in a 24-well plate using Lipofectamine RNAiMAX Transfection Reagent (Thermo Fisher Scientific, USA, 13778075).

ARA-elncRNA1 overexpression experiments were performed on LNCaP and 22Rv1 cells using pcDNA3.1-ARA-elncRNA1 plasmid and pcDNA3.1 as a control. For transfection Turbofect (Thermo Fisher Scientific, USA, R0531) was used according to the manufacturer’s protocol.

### RNA extraction and quantitative real-time RT-PCR (RT-qPCR)

RNA was isolated using the HiPure Total RNA Kit (Magen Biotechnology Co., China) according to the manufacturer’s protocol. Residual DNA was removed using DNase I (Magen Biotechnology Co., China). The first strand of cDNA was synthesized from 1 µg of total RNA using the Maxima H Minus Reverse Transcriptase (Thermo Fisher Scientific, USA) according to the manufacturer’s protocol, in a final reaction volume of 20 µL. The resulting cDNA mixture was diluted with 20 µL of nuclease-free water, and 2 µL of the diluted cDNA was used for each technical replicate in qPCR analysis. qPCR was performed using a 5X qPCRmix-HS SYBR (Evrogen, Russia).

### Cell fractionation and RNA extraction

Protocol from [43] was used for cell fractionation with minor modifications. Briefly, 10⁷ R1881 treated LNCaP cells were harvested using trypsin, washed twice with PBS, and centrifuged at 200 × g for 5 minutes at 4 °C. The cell pellet was resuspended in ice-cold Igepal buffer (10 mM Tris, pH 7.4; 150 mM NaCl; 0.15% Igepal CA-630) and incubated on ice for 5 minutes. A 20 µL aliquot of the lysate (5%) was saved as the input for total RNA extraction using TRIzol (Thermo Fisher Scientific, USA). The lysate was overlaid onto sucrose buffer (10 mM Tris, pH 7.4; 150 mM NaCl; 24% sucrose) and centrifuged at 3500 × g for 10 minutes at 4 °C. The supernatant was collected and saved for further analysis. A 100 µL aliquot of the supernatant was used for cytoplasmic RNA extraction using 500 µL of TRIzol (Thermo Fisher Scientific, USA).

The nuclear pellet was resuspended in 250 µL of glycerol buffer (20 mM Tris, pH 7.4; 75 mM NaCl; 0.5 mM EDTA; 50% glycerol) and 250 µL of urea buffer (10 mM Tris, pH 7.4; 1 M urea; 0.3 M NaCl; 7.5 mM MgCl₂; 0.2 mM EDTA; 1% Igepal CA-630). The mixture was vortexed for 4 seconds and incubated on ice for 2 minutes, followed by centrifugation at 13000 × g for 2 minutes. The supernatant, containing the nucleoplasmic fraction, was collected and saved. A 100 µL aliquot of this supernatant was used for nucleoplasmic RNA extraction with TRIzol (Thermo Fisher Scientific, USA). The chromatin pellet was dissolved in 500 µL of TRIzol for chromatin RNA extraction.

Input, cytoplasmic, nucleoplasmic, and chromatin RNA were reverse-transcribed into cDNA using Maxima H Minus Reverse Transcriptase (Thermo Fisher Scientific, USA) and analyzed by RT-qPCR. All buffers were supplemented with Complete Protease Inhibitor Cocktail (Merck, 11697498001) and 40 U/mL RNase inhibitor SUPERase•In.

### Nuclear and cytoplasmic extracts

Cells were harvested using trypsin, washed twice with ice-cold PBS, and centrifuged at 200 × g for 5 minutes at 4 °C. The pellet was gently resuspended in 500 µL of hypotonic buffer (20 mM Tris-HCl, pH 7.4; 10 mM NaCl; 3 mM MgCl₂; and protease inhibitors (Merck, 11697498001)) and incubated on ice for 15 minutes. Next, 25 µL of 10% NP-40 was added to the lysate, and the mixture was vortexed for 10 seconds at the highest setting. The homogenate was centrifuged at 3,000 rpm for 10 minutes at 4 °C. The supernatant, representing the cytoplasmic fraction, was collected and saved for further analysis (−80 °C).

### RNA immunoprecipitation (RIP)

5×10⁶ LNCaP or 22Rv1 cells treated with R1881 were harvested using trypsin, washed twice with ice-cold PBS, and separated into nucleolar and cytoplasmic fractions as described previously. For classical RIP, the nuclear pellet was lysed in 500 µL of RIP buffer (150 mM KCl, 25 mM Tris pH 7.4, 5 mM EDTA, 0.5% NP-40, 0.5 mM DTT, 40 U/mL RNase inhibitor SUPERase•In (Thermo Fisher Scientific, USA), and protease inhibitors (Merck, 11697498001)). A 50 µL aliquot of the lysate was saved as the input control. The lysate was incubated overnight with one of the following antibodies: 5 µL of anti-AR (Cell Signaling, 5153), 10 µL of anti-YY1 (Cell Signaling, 46395), 5 µL of anti-EZH2 (AbClonal, A5743), anti-SUZ12 (AbClonal, A4348), anti-H3 (Cell Signaling, 9715), or 10 µL of IgG (Cell Signaling, 2729). RNA-protein complexes were precipitated overnight at 10 °C using 40 uL of Pierce Protein A/G Magnetic Beads (Thermo Fisher Scientific, USA) and washed three times with RIP buffer. RNA was extracted using TRIzol reagent (Thermo Fisher Scientific, USA). cDNA was synthesized using Maxima H Minus Reverse Transcriptase (Thermo Fisher Scientific, USA) from even volumes of each RNA sample, and samples were analyzed by RT-qPCR.

For RIP of AR in the cytoplasmic fraction, the cytoplasmic fraction in hypotonic buffer was supplemented with KCl, EDTA, DTT, and SUPERase•In to match the RIP buffer composition. The RIP experiment was then performed as described above.

### Chromatin immunoprecipitation (ChIP)

Chromatin immunoprecipitation (ChIP) was performed using the SimpleChIP Plus Enzymatic Chromatin IP Kit (Cell Signaling, #9005) according to the manufacturer’s protocol. Briefly, 4 × 10^6^ R1881 treated cells were fixed with 1% formaldehyde, and cell nuclei were isolated. Chromatin was then fragmented using a micrococcal nuclease. The fragmented chromatin was incubated overnight with one of the following antibodies: 5 µL of anti-AR (Cell Signaling, 5153), 10 µL of anti-YY1 (Cell Signaling, 46395), 5 µL of anti-EZH2 (AbClonal, A5743), or 10 µL of IgG (Cell Signaling, 2729). A 2% aliquot of the chromatin fraction was collected prior to antibody addition and used as the Input control. Antibody-bound DNA was precipitated using magnetic beads. DNA was purified from the protein complexes and the Input fraction, and 2 µL of each sample was used for real-time qPCR analysis. qPCR was performed using a 5X qPCRmix-HS SYBR (Evrogen, Russia). Percent of input was calculated using the formula:

Percent Input = 2% x 2^(C[T]^ ^2%Input^ ^Sample-C[T]^ ^IP^ ^Sample)^, where C[T] = CT = Threshold cycle of PCR reaction. Primer sequences are listed in Table S4.

### Chromatin Isolation by RNA Purification (ChIRP)

The protocol from Chu et al. [44] was used with minor modifications. Briefly, four biotin-tagged DNA oligonucleotides complementary to the ARA-elncRNA1 sequence (Table S4) were designed using Biosearch Technologies’ Stellaris FISH Probe Designer. As a control, four oligonucleotides targeting the LAC-Z operon from a previous study [45] were used (Table S4). All biotin-tagged DNA oligos were purchased from Syntol (Russia).

A total of 3 × 10⁷ R1881-treated LNCaP cells were harvested using trypsin and resuspended in 20 mL of DMEM containing 1% formaldehyde. The cells were incubated at room temperature for 10 minutes with gentle rotation. Crosslinking was quenched by adding 2 mL of 1.25 M glycine and incubating with continued rotation at room temperature. The cell pellet was collected by centrifugation at 2000 x g for 5 minutes at 4 °C.

Cells were lysed in Lysis Buffer (50 mM Tris-HCl, pH 7.0, 10 mM EDTA, 1% SDS, protease inhibitor cocktail (PIC), and SUPERase•In) and sonicated for 20 minutes (Φ2 tip, 100 W, 1 s impulse, 1s rest, Ultrasonic Processor Scientz-IID, Scientz Bio, China). Chromatin was cleared by centrifugation at 16000 × g at 4 °C and divided into two parts. A 10 µL aliquot was taken as the input sample.

To one volume of chromatin, two volumes of hybridization buffer (750 mM NaCl, 50 mM Tris-HCl, pH 7.0, 1 mM EDTA, 1% SDS, 15% formamide, PIC, SUPERase•In) and 100 pmol of the oligo mix were added. The mixture was incubated at 37 °C with rotation overnight. Subsequently, 100 µL of Dynabeads MyOne Streptavidin C1 (Thermo Fisher Scientific, USA) was added to each sample and incubated with rotation at 37 °C for 30 minutes.

The beads were separated using a magnet stand and washed five times. They were then incubated in elution buffer (50 mM NaHCO₃, 1% SDS) containing 10 µL of RNase A (10 mg/mL, Evrogen, Russia) at 37 °C for 30 minutes with shaking. The input sample was diluted with 140 µL of elution buffer.

Both the ChIRP and input samples were incubated with 15 µL of proteinase K at 50 °C for 45 minutes with shaking. DNA was extracted using the ChIP DNA Clean & Concentrator kit (D5201, Zymo Research, USA) and eluted in a final volume of 30 µL. A 2 µL aliquot of the sample was used for qPCR analysis.

### Electrophoretic Mobility Shift Assays (EMSA)

#### RNA EMSA (rEMSA)

The protocol was adapted from Ream et al. [46]. Briefly, a 2.5% agarose gel in 0.5× TBE buffer (45 mM Tris, 45 mM boric acid) was used for electrophoretic separation of protein:RNA complexes. RNA oligonucleotides 5’-CAACUUCUCCUCUCU-Cy3-3’ and 5’-CAACUUCUCCUCUCU-3’, each containing an RNA-binding motif, were purchased from Syntol (Russia).

For the binding assay, 1 pmol of the Cy3-tagged RNA oligonucleotide was incubated with 2, 4, or 6 µg of total 22Rv1 R1881 treated cell lysate in Binding Buffer (5×: 200 mM Tris pH 8.0, 150 mM KCl, 5 mM MgCl₂, 0.05% (w/v) NP-40, 5 mM DTT, 25% glycerol) supplemented with 1 µg of total *E. coli* RNA as a nonspecific competitor, 10 µg/mL BSA, and a protease inhibitor cocktail (Merck, 11697498001). The reaction was incubated for 1 hour in the dark at room temperature. To test binding specificity, a 50-fold excess of untagged competitor RNA was added to one of the samples. To verify AR binding to the RNA oligonucleotide, 0.5 µL of anti-AR antibody (Cell Signaling, 5153) was added to the samples. The gel was run at 80 V until the control dye was 1–2 cm from the bottom, at 10 °C. Protein:RNA complexes were visualized using a ChemiDoc MP Imaging system (BioRad, USA).

#### DNA:DNA:RNA Triplex Formation Analysis

The protocol from Leisegang et al. [26] was used with minor modifications. Predicted DNA and RNA oligonucleotides for triplex formation were purchased from Syntol (Russia): DBD_RNA: 5’-AAAAAAAAAAAAAAAAAAAACUCUA-Cy5-3’, DNA1_F:

5’-AAAACAGTAATAATACATTTGGGGA-3’, DNA2_R:

5’-TCCCCAAATGTATTATTACTGTTTT-3’. The DNA-binding site is located in the *CEBPB* gene promoter.

For triplex formation, complementary DNA oligonucleotides were first incubated at 95 °C for 5 minutes in hybridization buffer (25 mM HEPES pH 7.4, 50 mM NaCl, 10 mM MgCl₂) and then gradually cooled to 25 °C (1 °C per minute). Next, 4 pmol of the tagged RNA oligonucleotide was mixed with varying amounts of dsDNA oligonucleotide (0, 2, 4, 6, or 16 pmol) and incubated at 60 °C for 1 hour, followed by cooling to 25 °C in a final volume of 10 µL. After incubation, 5 µL of 25% glycerol was added to each sample.

A native PAGE gel was prepared using polyacrylamide gel (containing 12% (w/v) acrylamide, and 3.0% (w/w) bisacrylamide) and 0.5× TBE buffer (45 mM Tris, 45 mM boric acid). Electrophoresis was performed at a constant voltage of 90 V for 1.5 hours. DNA:DNA:RNA triplexes were visualized using a ChemiDoc MP Imaging system (BioRad, USA).

### Cell lysis and western blot analysis

Cells from one well of a 12-well plate were lysed in 50 µL of 1× Cell Lysis Buffer (Cell Signaling, 9803) supplemented with 1 mM PMSF (PMSF-RO, Roche). The lysate was centrifuged at 4 °C and 16,000 × g for 10 minutes. The supernatant was collected and stored at −80 °C for further analysis. Protein concentration was determined using a BCA Protein Assay Kit (Thermo Fisher Scientific, USA).

For Western blot analysis, 40 µg of each sample was used. Briefly, 4x Laemmli buffer was added to each sample, followed by boiling at 95 °C for 5 minutes. Electrophoretic separation of proteins was performed using a Bio-Rad Mini Protean chamber. Proteins were transferred to a PVDF membrane (Bio-Rad, USA) using a Mini Trans-Blot Cell (Bio-Rad, USA). The membrane was blocked in PBST containing 5% nonfat dry milk and then incubated with primary antibodies against TUB (1:1000, PAB870Hu012, Cloud-Clone Corp., USA) or AR (1:1000, 5153, Cell Signaling Technology, USA) overnight at 4 °C. After washing with PBST, the membrane was incubated with horseradish peroxidase (HRP)-conjugated secondary antibodies (31460, Thermo Fisher Scientific, USA) for 1 hour at room temperature. Protein bands were visualized using Pierce ECL Western Blotting Substrate (Thermo Fisher Scientific, USA) on a ChemiDoc MP Imaging system (Bio-Rad, USA).

### Quantitative chromosome conformation capture (3C) Assay

3C assays were performed as previously described [47]. Briefly, LNCaP cells in 10 cm dishes were transfected with siRNA targeting ARA-elncRNA1 or with siRNA targeting LUC as a control. Similarly, 22Rv1 cells in 10 cm dishes were transfected with pcDNA3.1-ARA-elncRNA1 or with pcDNA3.1 as a control. The following day, the medium was replaced with a full growing medium DMEM containing 10 uM R1881. After treatment, the cells were crosslinked, and chromatin was purified.

The crosslinked chromatin was digested overnight with the MboI restriction enzyme (Fermentas, USA) following DNA ligation with T4 DNA ligase (Thermo Fisher Scientific, USA). After crosslink reversal, the DNA was purified by phenol/chloroform extraction. A total of 50 ng of DNA was used for qPCR analysis.

Ligation frequencies for each sample were normalized to the qPCR signal from a gene desert region on chromosome 12. Primers used for qPCR are listed in the Table S4.

### Colony formation assay

For each transfected or transduced cell line, 5,000 cells per well of 6 well plate were seeded. All samples were incubated in a CO_2_ incubator for 2 weeks. Cell colonies were fixed with 7 parts of methanol and 1 part of glacial acetic acid. Samples were stained with 0.5% (w/v) trypan blue in mQ for 30 minutes and washed 3–5 times with PBS. To quantify the number of cell colonies and average area of a colony ImageJ software was used [48].

### Cell proliferation assay

Cells were seeded at a density of 70,000 cells per well in a 24-well plate and transfected with siRNA or DNA plasmid if needed. Cell proliferation was measured 24h after treatment using the Celena X High Content Imaging System (Logos Biosystems, South Korea) over a 48-hour period. Images were acquired every 2 hours at 4× or 10× magnifications.

To quantify cell proliferation, the total area occupied by cells in each image was analyzed using CellProfiler software [49]. Three biological replicates were performed for each condition.

## Statistical analysis

Results were analyzed using two-tailed unpaired Student’s t-test and Fisher’s exact test. P-value < 0.05 was considered statistically significant.

## Supplementary information

### Supplementary figures

**Figure S1.**
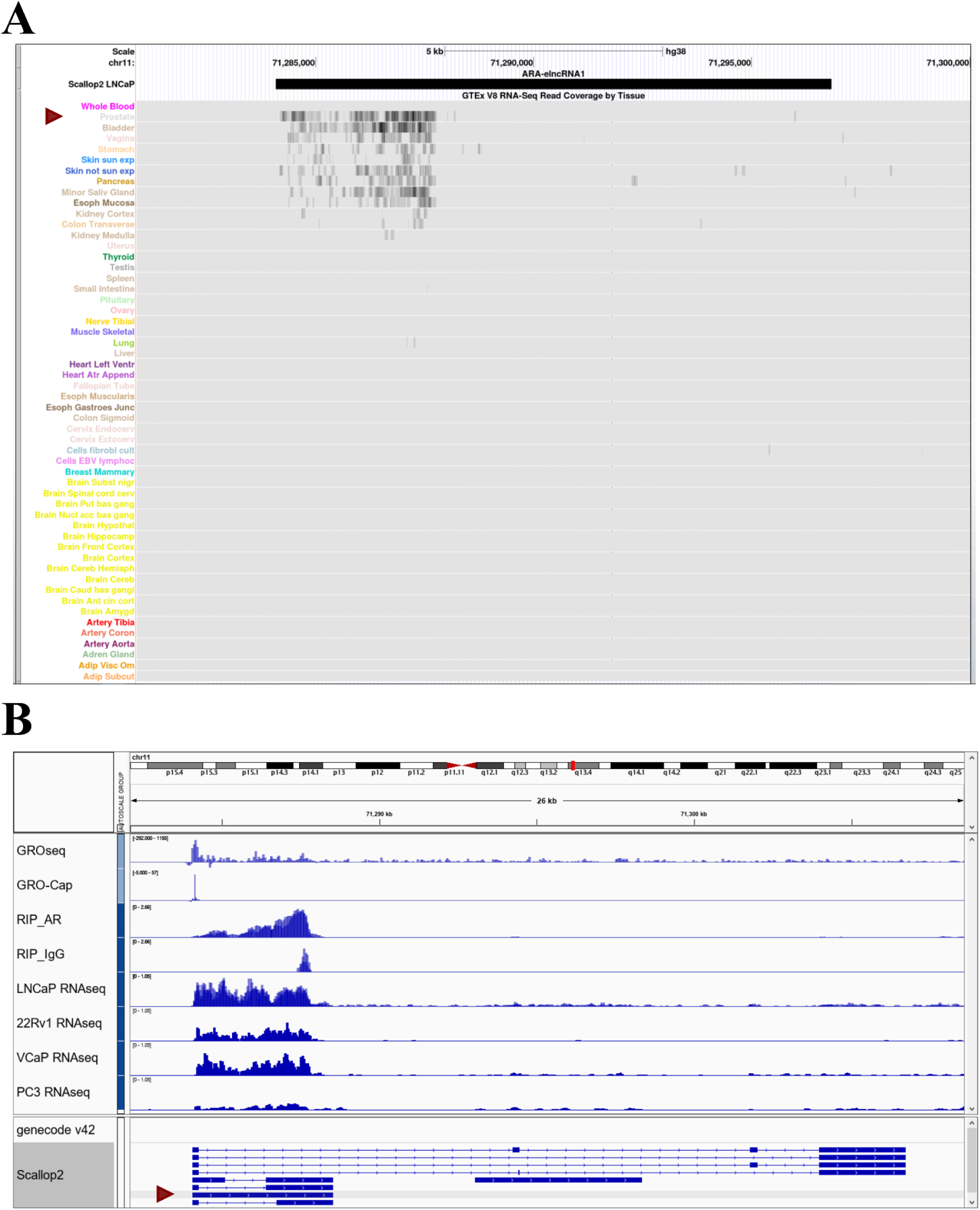
S1A. Snapshot of UCSC Genome Browser with GTEx RNAseq data tracks. Red triangle indicates RNAseq data track for prostate tissue. S1B. Snapshot of The Integrative Genomics Viewer with GENCODE v42 gene annotation, scallop2 *de novo* gene assembly track, RNAseq of LNCaP, 22Rv1, VCaP and PC3 cells, AR and IgG RIP-seq data track for LNCaP, GRO-seq and GRO-Cap data tracks for LNCaP. Red triangle indicates a 4kb length monoexonic transcript of gene.56338.3 which we designate as ARA-elncRNA1.

**Figure S2.**
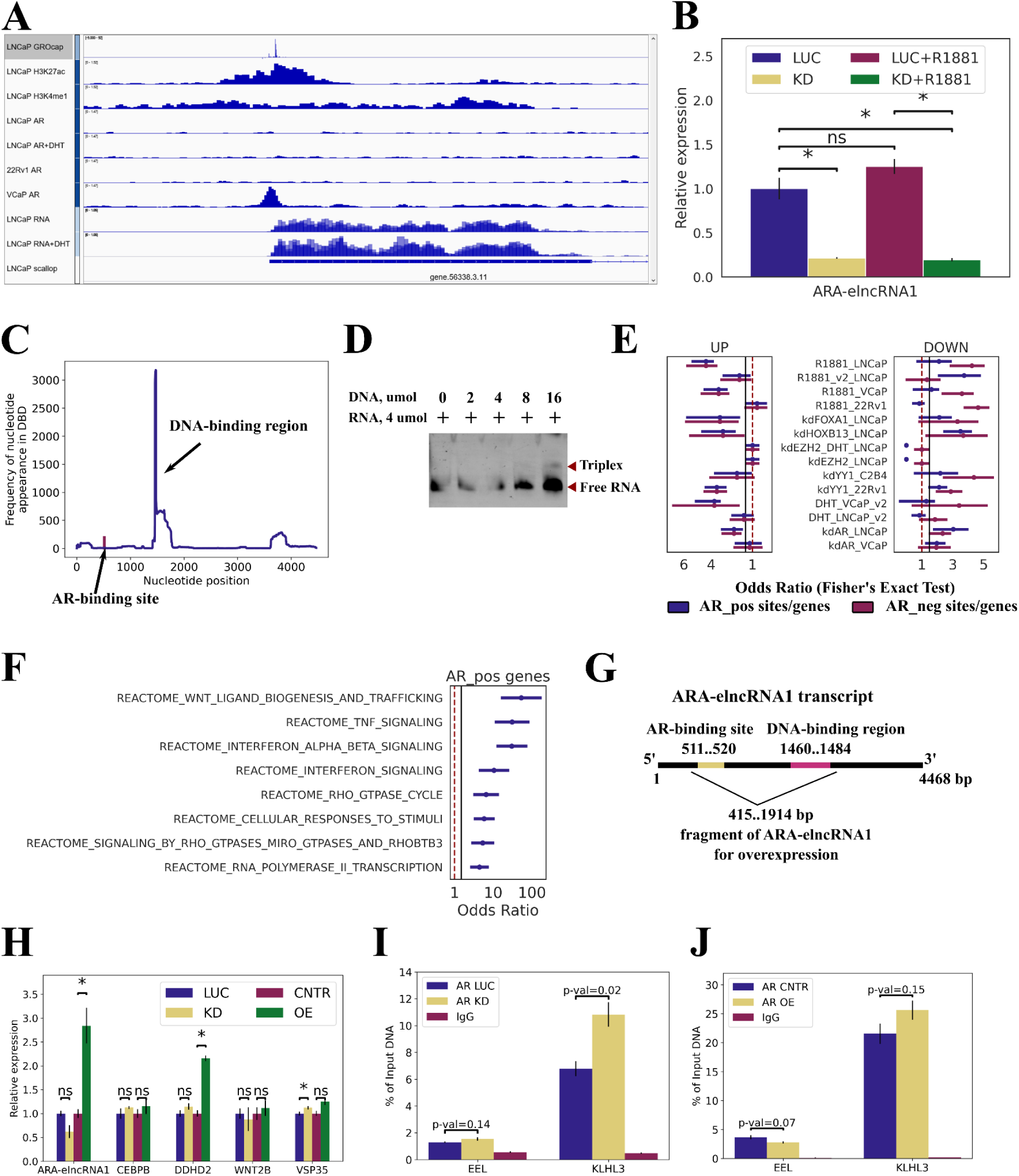
S2A. Snapshot of the IGV genome browser showing the scallop2 *de novo* gene assembly track, RNA-seq data for LNCaP and LNCaP+DHT, GRO-Cap data track, H3K27ac, H3K4me1, AR, AR+DHT ChIP-seq for LNCaP, and AR ChIP-seq for 22Rv1 and VCaP cells. S2B. Relative normalized expression of ARA-elncRNA1 following ARA-elncRNA1 knockdown (KD) and R1881 treatment in LNCaP cells. “*” indicates p-value < 0.05; “ns” indicates non-significant (two-tailed unpaired Student’s t-test). Results are presented as mean ± SEM from three replicates. LUC - control siRNA duplex to firefly luciferase. S2C. Frequency of nucleotide occurrence in the DNA-binding region predicted by TriplexAligner. The region 1460–1484 is designated as the DNA-binding domain (DBD) of ARA-elncRNA1. S2D. Electrophoretic mobility shift assay (EMSA) of the *CEBPB* DNA duplex and the DNA-binding domain of ARA-elncRNA1. S2E. Enrichment analysis of AR_pos and AR_neg genes with up-and down-regulated genes following R1881/DHT treatment or knockdown (KD) of corresponding genes in the listed cell lines. Odds ratios with 95% confidence intervals are shown (Fisher’s exact test). S2F. Enrichment analysis of AR_pos genes with Reactome pathways. Selected pathways are shown. Odds ratios with 95% confidence intervals are shown (Fisher’s exact test). S2G. Schematic overview of the 415–1914 region of ARA-elncRNA1, containing the AR-binding site (511–520) and the DNA-binding region (1460–1484), overexpressed in LNCaP or 22Rv1 cells. S2H-I. AR occupancy of two AR_neg sites in the *ELL* and *KLHL3* genes, measured by ChIP-qPCR in LNCaP cells under ARA-elncRNA1 KD (H) and in 22Rv1 cells under ARA-elncRNA1 overexpression (OE) (I). “*” indicates p-value < 0.05; “ns” indicates non-significant (two-tailed unpaired Student’s t-test). Results are presented as mean ± SEM from three replicates. LUC - control siRNA duplex to firefly luciferase. S2J. Relative normalized expression of ARA-elncRNA1, *DDHD2*, *VPS35*, *CEBPB*, and *WNT2A* genes following ARA-elncRNA1 KD or OE in 22Rv1 cells. “*” indicates p-value < 0.05; “ns” indicates non-significant (two-tailed unpaired Student’s t-test). Results are presented as mean ± SEM from three replicates. LUC - control siRNA duplex to firefly luciferase.

**Figure S3.**
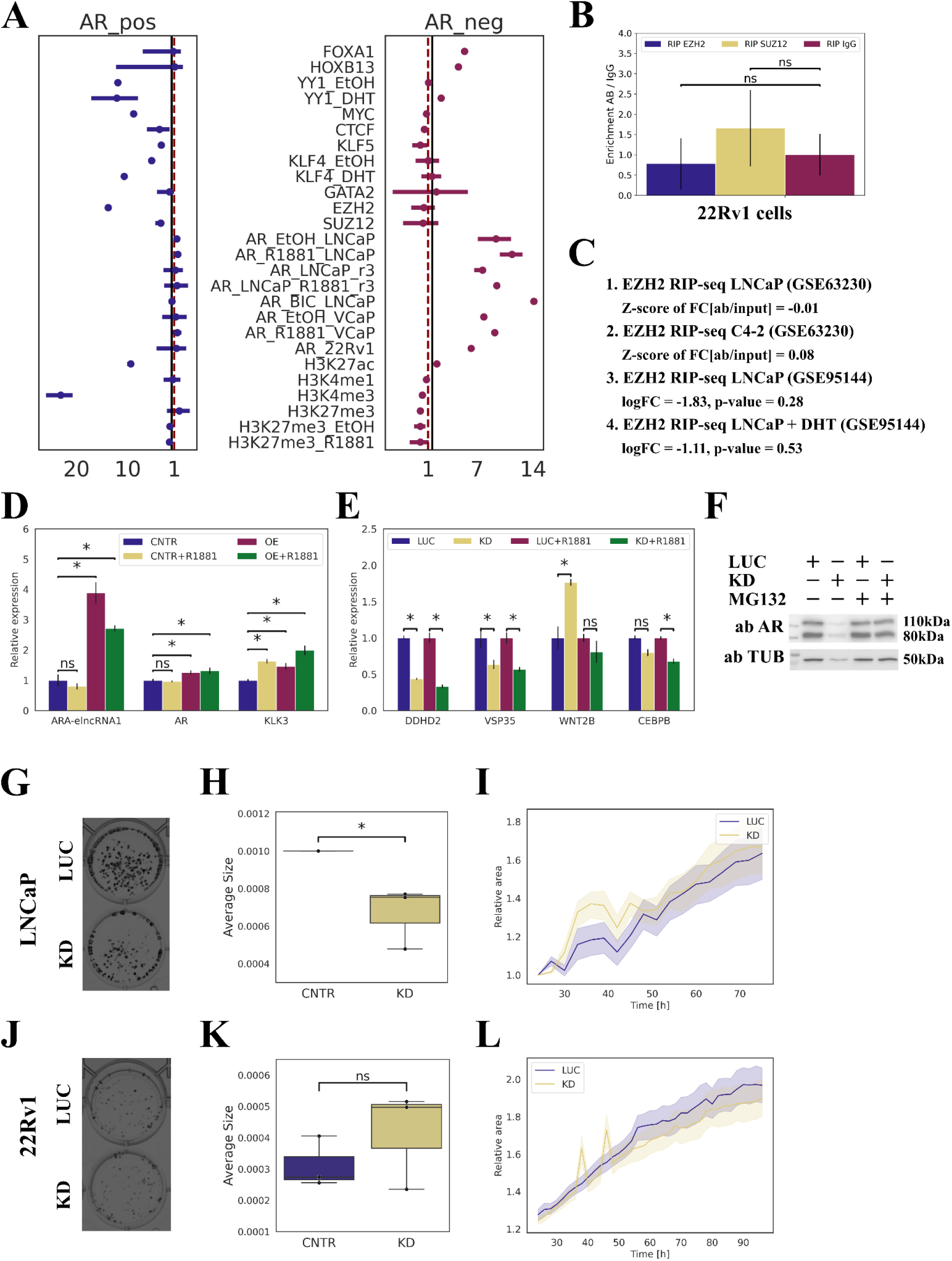
S3A. Enrichment of AR_pos and AR_neg sites with ChIP-seq peaks for different chromatin-associated proteins. Enrichment was measured using Fisher’s exact test. Sites were considered enriched if the Odds Ratio (OR) > 1.5 and FDR < 0.05. Odds ratios with 95% confidence intervals are shown. S3B. RIP results confirming the absence of interaction between ARA-elncRNA1 and EZH2 or SUZ12 in the nuclear fraction of 22Rv1 cells. RT-qPCR was used to assess the enrichment of ARA-elncRNA1 in the anti-EZH2 and anti-SUZ12 antibody fractions compared to IgG. “ns” indicates p-value > 0.05 (two-tailed unpaired Student’s t-test). Results are presented as mean ± SEM from three replicates. S3C. EZH2 RIP-seq analysis data for ARA-elncRNA1. Z-scores of Fold Change [antibody/input] are shown for experiments with one technical replicate (1, 2). LogFC and p-values from DESeq2 analysis are shown for experiments with three replicates (3, 4). S3D. Relative normalized expression of *AR* and *KLK3* genes following ARA-elncRNA1 overexpression (OE) and R1881 treatment in LNCaP cells. “*” indicates p-value < 0.05; “ns” indicates non-significant (two-tailed unpaired Student’s t-test). Results are presented as mean ± SEM from three replicates. S3E. Relative normalized expression of ARA-elncRNA1, *DDHD2*, *VPS35*, *CEBPB*, and *WNT2A* genes following ARA-elncRNA1 knockdown (KD) and R1881 treatment in LNCaP cells. “*” indicates p-value < 0.05; “ns” indicates non-significant (two-tailed unpaired Student’s t-test). Results are presented as mean ± SEM from three replicates. LUC - control siRNA duplex to firefly luciferase. S3F. Western blot analysis of 22Rv1 cells under ARA-elncRNA1 KD treated with MG132 (12.5 µM) for 4.5 hours (C), using anti-AR or anti-TUB antibodies. Numbers indicate relative abundance of AR compared to the control (LUC). LUC - control siRNA duplex to firefly luciferase. S3G-H. Colony formation assay of LNCaP cells under ARA-elncRNA1 KD. Representative images of colonies in wells are shown in panel G. Box plots represent the average size of a colony (H). Each experiment was performed in three biological replicates. “*” indicates p-value < 0.05; “ns” indicates non-significant (two-tailed unpaired Student’s t-test). S3I. Proliferation assay of LNCaP-mCherry cells under ARA-elncRNA1 KD. The line plot shows the relative area occupied by cells, measured using the Celena X High Content Imaging System over 48–100 hours. Images were acquired every 2–3 hours. The 95% confidence interval is shown as error bars. Each experiment was performed in three biological replicates. LUC - control siRNA duplex to firefly luciferase. S3J-K. Colony formation assay of 22Rv1 cells under ARA-elncRNA1 KD. Representative images of colonies in wells are shown in panel J. Box plots represent the average size of a colony (K). Each experiment was performed in three biological replicates. “*” indicates p-value < 0.05; “ns” indicates non-significant (two-tailed unpaired Student’s t-test). LUC - control siRNA duplex to firefly luciferase. S3L. Proliferation assay of 22Rv1-mCherry cells under ARA-elncRNA1 KD. The line plot shows the relative area occupied by cells, measured using the Celena X High Content Imaging System over 48–100 hours. Images were acquired every 2–3 hours. The 95% confidence interval is shown as error bars. Each experiment was performed in three biological replicates.

## Supplementary tables

Table S1. List of all datasets used in the current study.

Table S2. List of 1840 newly identified transcribed regions and their characteristics.

Table S3. List of regions with AR occupancy associated with ARA-elncRNA1 expression.

Table S4. List of all oligonucleotides used in the study.

## Supplementary files

File S1. GTF file containing *de novo* assembled transcripts for the LNCaP cell line.

File S2. FASTA file containing the nucleotide sequence of ARA-elncRNA1.

## Notes

### Competing Interest Statement

The authors have declared no competing interest.

### Summary of Updates

The abstract was missing its first paragraph. Necessary improvements have been added to it.

